# Differential temperature adaptation mechanisms in the High Arctic-adapted *Cerastium regelii* Ostenf. and the widespread *Stellaria longipes* Goldie

**DOI:** 10.1101/2025.03.15.643465

**Authors:** Sarah L. Lane, Lauren A E Erland

**Affiliations:** Agriculture, University of the Fraser Valley, Stó:lō Temexw, Abbotsford, BC, Canada, V2S 7M8

**Keywords:** chickweed, climate change, ecological niche model, long stalk starwort, metabolomics, Nunavut, plant environment resilience, temperature stress

## Abstract

- Climate change impacts arctic latitudes more acutely than other latitudes, resulting in arctic shrubification. How individual species in these climes respond to warming temperature is poorly understood. Understanding species resiliency to climate change will help us conserve plant species at risk.
- We performed a survey of plants in a permafrost anomaly in Resolute (Qausuittuq), Nunavut, Canada. Two identified species, *Stellaria longipes* Goldie and *Cerastium regelii* Ostenf., were investigated through modelled niche suitability under future climate scenarios, phenological analysis, and *in vitro* warming experiments to investigate growth and phytochemical profiles.
- 10 species including *Stellaria longipes* and *Cerastium regelii* were identified in the anomaly. Predicted niche suitability increased under SSP126 for *C. regelii*, with compressed and later flowering period since 1850*. In vitro, S. longipes* maximized growth at 24 °C with greater abundance of cytokinins than *C. regelii*, which increased growth at 28 °C.
- *Stellaria longipes* is self-limiting at higher temperatures, and is less temperature-dependent for its success, while *C. regelii* is more affected by warming temperatures, showing increases in growth and predicted niche suitability. Our work increases understanding of plant resiliency and vulnerability in Canada’s High Arctic, and sheds light on the biology of an understudied arctic specialist in *C. regelii*.

## Introduction

As climate change has shifted to become our global reality, high latitude and high altitude regions are experiencing the effects of these changes more acutely than other global regions (Rantanen *et al*., 2022). Inuit Nunangat is a distinct geopolitical region, with Inuit self-governance, which is also referred to as Canada’s High Arctic. While significant knowledge has been generated surrounding predicted impacts of climate change on terrestrial and marine ecosystems in this region, relatively little is still understood about the plants from a conservation perspective. For example, a review of the assessment status of medicinal plants in Canada in 2022 found that only 22 % of plant species have been assessed in Nunavut for a conservation ranking by NatureServe and related conservation data centres (Erland *et al*., 2022). Much of the available research has focused on woody plants examining the concept of shrubification of Arctic and tundra environments globally. Under these predicted future environments woody shrubs, including willows and perennial berry plants, are anticipated to grow taller with increasing growing degree days in their environments (Myers-Smith *et al*., 2011; Kodl *et al*., 2024). As primary producers, plants form the basis of ecosystems. Understanding how plants will respond to future environment scenarios can help to better predict and understand future environment ecosystem interactions.

Plant vulnerability and resiliency requires understanding how plants perceive and respond to environmental cues. Recent literature has proposed that vulnerability of plants to climate change is determined by three key factors: (a) exposure, i.e. extrinsic factors, (b) sensitivity, essentially the degree to which the plant is affected by the extrinsic factors, and (c) adaptive capacity, the ability for a plant to respond *in situ* to a changing climate and persist (Dawson *et al*., 2011; Foden *et al*., 2013). The first two factors have been widely investigated and the NatureServe Climate Change Vulnerability Index provides a foundation for identification of generalized adaptive traits, however, there is still a general lack of information on many of the traits mediating adaptive capacity. One of the ways that plants can interact with the world is through the production of specialized phytochemicals. Plant growth regulators or phytohormones are a set of phytochemicals produced by plants, which enable plants to perceive environmental changes and respond by redirecting growth, detoxifying stress metabolites and stabilizing physiological processes (Erland *et al*., 2017).

In 2019 we conducted a botanical survey and collection in Resolute (Qausuittuq) Nunavut, which included 83 vouchers and 74 seed collections, with 46 vouchers submitted to the BOLD database and the Canadian Museum of Nature Herbarium. As part of this effort, we worked with local Inuk guide, Brandy Iqaluk, and we identified an area of interest, which was a permafrost anomaly with increased temperature and associated modified plant community composition. This site unlike surrounding permafrost had an abundance of herbaceous species present. We hypothesized that this region could serve as a model for how the plant community composition could shift under future climate scenarios. This anomaly was dominated by the presence of two species in the family Caryophyllaceae which were present, but not abundant, in the surrounding areas. Therefore, our overall objective was to integrate data from field survey, ecological niche modelling, in vitro temperature experiments and metabolomics to predict and understand the differential impacts of environmental warming on two related herbaceous species growing in the Canadian High Arctic: *Cerastium regelii* Ostenf. and *Stellaria longipes* Goldie.

*Stellaria longipes*, longstalk starwort or Goldie’s starwort is a small herbaceous member of the Caryophyllaceae family, first identified in the woods of Ontario. It has a broad circumboreal distribution and has been found to grow in diverse ecosystem types including both arctic and alpine environments, though some debate exists in the literature about the potential existence of several ecotypes, species or subspecies regionally, with the species being referred to as a species complex (Porsild *et al*., 1980; Chinnappa *et al*., 2005; Sharples, 2019; ‘Stellaria longipes s. lat.’). Extensive previous work has established *S. longipes* as a remarkably phenotypically and morphologically plastic species, which likely contributes significantly to this confusion. Alpine and arctic specimens tend to have compact or dwarf phenotypes (referred to as alpine ecotypes), while plants growing in lower elevation and altitude locations including prairie ecosystems are much taller with longer internodal lengths (prairie ecotype). While initially hypothesized to be an environmentally sensitive species, as it is very responsive to temperature and photoperiod cues, *S. longipes* has been found to be remarkably resilient through in field experimentation (Macdonald *et al*., 1986; Emery *et al*., 1994; Maillette *et al*., 2000; Li *et al*., 2011) with the ecotypes have differential tolerance to stresses including drought stress (Sangtarash *et al*., 2009). The role of plant hormones including abscisic acid, gibberellins, ethylene, auxins and cytokinins in mediating these responses (Kathiresan *et al*., 1996; Kurepin *et al*., 2006, 2008; Li *et al*., 2011), especially stem elongation in response to changes in photoperiod and temperature (Macdonald *et al*., 1984, 1986; Thuy *et al*., 2009) (Macdonald *et al*., 1984, 1986; Thuy *et al*., 2009). Together this makes *S. longipes* an interesting and well characterized system for the study of environmental resilience (Chinnappa *et al*., 2005).

*Cerastium regelii* or Regel’s chickweed is a subarctic to arctic species found across the circumpolar region, with populations restricted primarily to above 70 °N (Heide *et al*., 1990). In contrast very little literature has been published on *C. regelii*, with for example a search for the species “*Cerastium regelii*” yielding only two documents on Web of Science, and only 242 observations globally recorded on iNaturalist (‘Regel’s chickweed’). Much of the available information is in the form of occurrence records and presence in botanical surveys e.g. (Hanssen, 1932; Fredskild, 1984; Matveyeva, 1994) or inclusion in phylogenetic studies e.g. (Yao *et al*., 2021; Liu *et al*., 2023). A study of plant community composition after glacial moraine retreat in Svalbard found that *C. regelii* had higher occurrence in sites with higher amounts of fine texture soil cover. Heide et al. 1990 describe photoperiod and flowering time requirements for this species, noting that while it is a short day plant, it grows exclusively in a region where it would experience long day lengths during the flowering season. The authors also identify *Cerastium jenisejense* (tundra chickweed) as a synonym and high temperature morphotype of *C. regelii* (Heide *et al*., 1990; Kew Science), though inclusion of the synonym in our Web of Science search does not expand the literature available. Lastly, a cyclopeptide deemed ‘regelin A’ has been isolated from *C. regelii* with the primary interest in cyclopeptides being as natural products (Ai-Qun *et al*., 2004; Zorzi *et al*., 2017).

We hypothesize that *S. longipes*, a generalist species with a pan-Canadian distribution would have higher temperature resilience as defined by enhanced growth under a wider temperature range, than *C. regelii*, and that this would be associated with modified phytohormone metabolism profiles. Guided by our in-field observations, we use three main approaches to investigate this hypothesis (1) in vitro germination studies, (2) phenology & ecological niche modelling and (3) untargeted hormonomics studies. We generate new information on the little studied arctic chickweed *C. regelii* and generate novel hypotheses for the role of phytohormones in environmental resilience.

## Materials and Methods

### Botanical Collections and Ecological Surveys

Field samples of growing aerial tissues of *Cerastium regelii* Ostenf. and *Stellaria longipes* Goldie were collected in July - August 2019 from sites just outside the Hamlet of Resolute, Qausuittuq, Nunavut, near 3 Mile Lake (74.77011, −95.09212). Samples were collected under Wildlife Research Permit #2019-047 granted by the Nunavut Department of Environment pursuant to the Wildlife Act. Working with local Inuk guide Brandy Iqaluk, we asked “Are there areas that you have noticed in which the plants are changing in recent years?”. An area of interest was identified for sampling due to its unusually high temperature compared to the surrounding areas and unique plant community composition. This site showed no signs of animal activity such as a lemming den or carcass which may explain the increased plant density and Ms. Iqaluk did not know of any previous use of the site which could account for, for example, a greater nutrient availability.

Thermal readings of the site were conducted using a FLIR ONE Pro thermal infrared camera and the FLIR ONE app (Teledyne FLIR, USA). The feature was divided into three zones for survey of the vegetation based on the differences in the plant community and “inside”, “middle” and “outside” region (Figure 1) which had distinct community composition. Detailed photos were taken of each of the regions. The plants present were identified in the field and density was counted based on the number of stems. While this does not provide an accurate representation of the number of individual plants due to the dense matte forming growth form of the chickweeds, it provides an approximate of the coverage stem density of the species.

**Figure 1.**
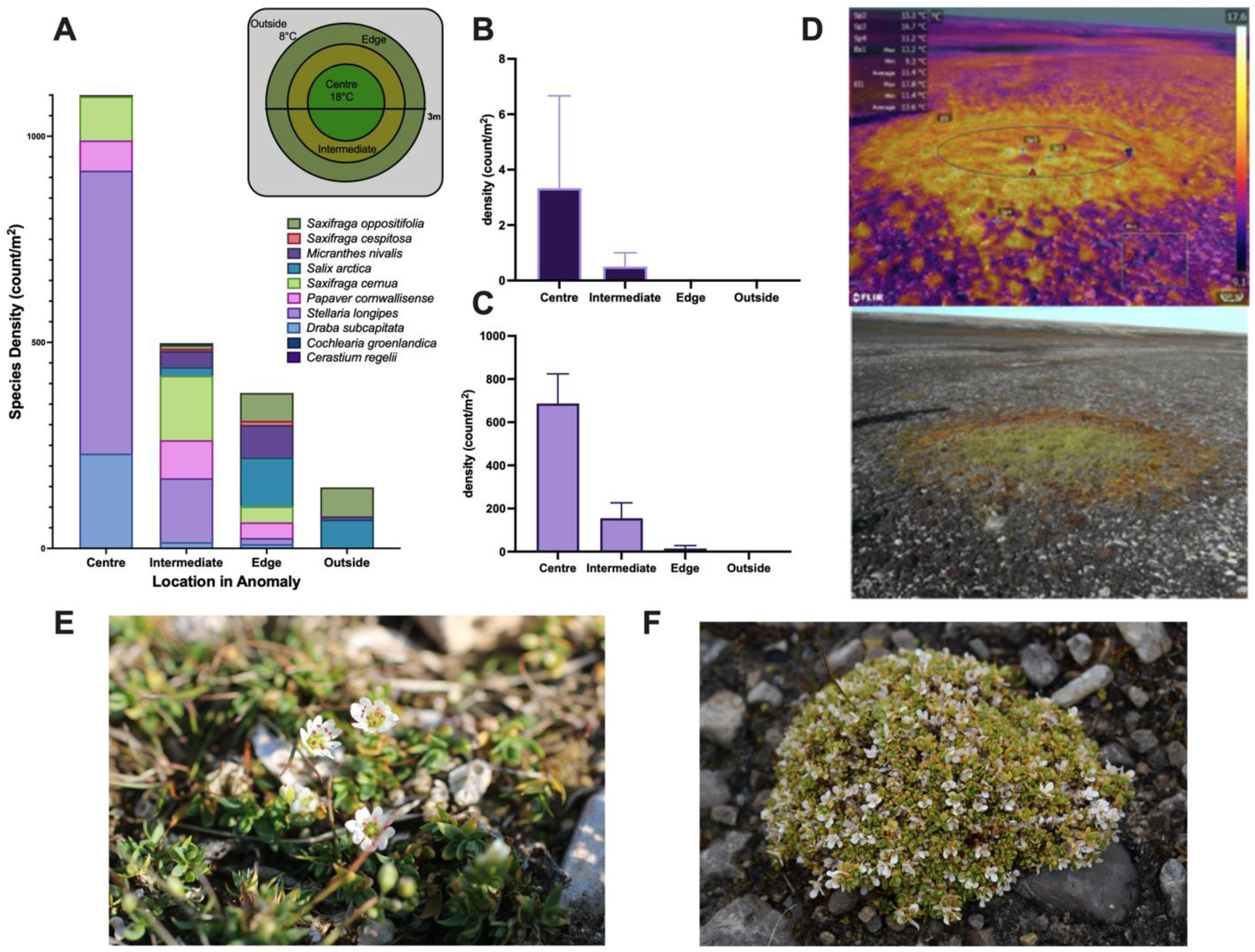
Plant community composition at the permafrost feature (a), density of (b) Cerastium regelii and (c) Stellaria longipes at the sites (d) thermal camera readings overlaid with RGB (bottom) image of the anomaly and field photos from Resolute (Qausuittuq), NU of: (e) S. longipes and (f) C. regelii. Error bars represent standard error.

For all species botanical vouchers were collected and were submitted for DNA barcoding and confirmation of botanical identification to the Herbarium at the Canadian Museum of Nature (BOLD Project ID: CCDB-25796).

Seed and aerial tissue samples were collected. Tissues were collected and transported in nutrient solution composed of one electrolyte tablet (Nuun Hydration Sport “Citrus Fruit”, NPN 80089152) in 500 mL of water with 1% Head and Shoulders shampoo containing zinc pyrithione (Classic, Proctor & Gamble), prior to establishment of cultures at the University of British Columbia, Okanagan. Seed samples were isolated from receptacle tissue and stored in envelopes at −20 °C.

### Establishment of an in vitro germplasm collection in controlled environments

Field-collected explants were transported from Nunavut to Kelowna, BC over 9 days. Upon arrival in the lab, explants were surface sterilized in a 10 % sodium hypochlorite (Chlorox ® Bleach) with 1 % Head and Shoulders shampoo for 15 minutes followed by three rinses with sterile distilled water. Explants were cultured onto Murashige and Skoog basal media (MSO) (Murashige & Skoog, 1962) containing full strength MS salts with Gamborg’s B5 Vitamins (Gamborg *et al*., 1968), 3% sucrose and 0.25 % phytagel, pH adjusted to 5.75 using NaOH. All media was sterilized by autoclave (Steris, Mentor, OH, USA; 121 °C at 18 psi for 20 min) prior to being poured across 100 mm x 15 mm petri dishes (Thermo Fisher Scientific, Canada). Sequential surface sterilization and re-subcultures of field-collected explants was necessary to establish axenic material for the germplasm collection and chemical studies. Cultures were maintained in a growth chamber at 21 °C under full spectrum white light (Sunblaster™ full spectrum daylight fluorescent bulbs; 40 μmol/m^2^/s), with a 16 h photoperiod. Once established, species were subcultured as needed when media was spent or tissues had fully filled the plate.

### Botanical Records, Species Distribution

Historical occurrence records for both species were downloaded from the Global Biodiversity Information Facility (GBIF):

*Cerastium regelii* (LAEE0038):https://doi.org/10.15468/dl.pcu6j5

*Stellaria longipes* (LAEE0051):https://doi.org/10.15468/dl.6uqxtz)

Data was cleaned to include only ‘SPECIES’ (and for *S. longipes* also ‘SUBSPECIES = longipes’) and country ‘CA’, duplicates were removed and the data was formatted in species with data (SWD) format. Occurrences recorded during the 2019 field collection trip were then added.

Changes in phenology in the historic record were screened. Occurrence data acquired from the GBIF used in ecological niche models was manually screened for the presence of flowers, flower buds and seeds. Data was grouped into the date ranges 1850-1950, 1951 – 1975, 1976-2000, 2000-2010 ad 2011-2019. The proportion of the observations with visible flowers or flower buds was calculated by month and plotted in Prism (GraphPad, v10, RRID:SCR_002798).

### Ecological Niche Modelling

Ecological niche modelling using the maximum entropy model (Maxent) was performed following procedures previously described (Hirabayashi *et al*., 2022) with modification. In brief, ENMEvaluate, ENMTools and ecospat R packages were used for variable selection and to optimize model settings (regularization multiplier and feature type) and variables retained for assessment are detailed in Table S1 (Muscarella *et al*., 2014; Kass *et al*., 2021). ENMEvaluate was run with spatial block partitions, testing regularization multiplier (RM, 1-5) and feature type (linear (L), quadratic (Q), hinge (H), LQ or LQH), with parallel = TRUE, algorithm = Maxent.jar, all other settings were left as default. Random background points were generated in QGIS (v3.16.14 long term release, QGIS.org, 2021) using the Database of Global Administration Areas (GADM) shapefile (Level 0) for Canada (www.gadm.org) as the extent. Optimal models were selected based on 10 % omission rate, and where similar choosing the model with the lower Akaike information criterion (AICc) value (Muscarella *et al*., 2014; Kass *et al*., 2021). Null models were constructed using the optimized parameters as an additional quality check to ensure empirical > null (Table S2).

Environmental variables including landcover (Tuanmu & Jetz, 2014), topography (Amatulli *et al*., 2018), edaphic properties (Poggio *et al*., 2021), habitat heterogeneity (Tuanmu & Jetz, 2015) and bioclimatic variables (Fick & Hijmans, 2017) were assessed and selected to avoid collinearity based on correlations as reported by the dataset authors original reports (Tuanmu & Jetz, 2014, 2015; Fick & Hijmans, 2017; Amatulli *et al*., 2018; Poggio *et al*., 2021) and as described in Hirabayashi et al. 2022 (Table S2). Soil and edaphic properties were downloaded from the SoilGrids2 Web Coverage Service (WCS mean @ 0-5 cm) as both species have relatively shallow root systems. Topographic, consensus landcover and habitat heterogeneity variables were downloaded from www.earthenv.org at 30 arc seconds resolution based on GMTED2010 and median aggregation where applicable (Amatulli *et al*., 2018). Consensus land cover variables were downloaded with DIScover (includes integration of Global Land Cover Characterization, GLCC v2.0) and included: regularly flooded vegetation, cultivated and managed vegetation, snow and ice cover (Tuanmu & Jetz, 2014). Habitat heterogeneity variables included: homogeneity, evenness, entropy, dissimilarity, and coefficient of variation (coeffvar) (Tuanmu & Jetz, 2015). Historic bioclimatic variables were downloaded from WorldClim v2.1, RRID:SCR_010244 at 30 arc second resolution. Future climate data (bioclimatic variables) were downloaded from www.worldclim.org for the Coupled Model Intercomparison Project (CMIP) 6 multi-model ensemble under the least and most extreme shared socioeconomic pathways (SSPs): 126 (lowest emissions, smallest increase in temperature) and 585 (highest emissions, largest increase in temperature) averages; for projections to 2041-2060 and 2061-2080 (referred to as 2060 and 2080). Three global climate models (GCMs): CNRM-CM6-1 (CMIP6.CMIP.CNRM-CERFACS.CNRM-CM6-1; Voldoire *et al*., 2019), MIROC6 (CMIP6.ScenarioMIP.MIROC.MIROC6; Shiogama *et al*., 2019) and MRI-ESM2-0 (CMIP6.CMIP.MRI.MRI-ESM2-0; Yukimoto *et al*., 2019) were selected based on best performance in GCMEval (gcm.met.no) using focus regions: Alaska/NW Canada and Canada/Greenland/Iceland with equal weights for skill evaluation (Parding *et al*., 2020; Shiogama *et al*., 2021). All variables were resampled to a resolution of 30 arc seconds in RStudio.

Maxent was run in dismo in R (RRID:SCR_001905) with RM = 4 for *C. regelii* and = 5 for *S. longipes*. All other parameters were as follows: feature class = linear quadratic hinge, autofeatures = FALSE, using maximum of 10,000 background points, maximum of 5,000 iterations, random seed = TRUE, allowing partial data using raster grids of selected environmental variables. Ten replicate cross-validation runs, withholding one fifth of the data as testing data were used for training and Maxent was then run using optimized settings for each species for historic and future climate scenarios. Receiver operator area under the curve (AUC) and true skill statistic (TSS) were calculated using the SDMtune R package using optimized Maxent settings for each species and withholding 20% of data as a testing set and using k-fold cross validation (Vignali *et al*., 2020). Scores are reported for training, testing and heldback datasets (Table S3). The output for the GCMs (CNRM-CM6-1, MIROC6 and MRI-ESM2-0) used under each SSP and at each timeframe were then averaged and the mean model output (cloglog) was visualized in QGIS.

### Temperature Response Experiments

Two node long cuttings were taken from *in vitro* cultured in established plants for each species, with four explants (pseudoreplicates) started per petri plate (12 °C, 24 °C, or 28 °C). Cultures were maintained under cool white fluorescent lights (∼40 µmol m^-2^ s^-1^, Sunblaster T4) with a 16 h photoperiod. Cultures grown for three weeks under test conditions at which point samples were destructively sampled and growth characteristics measured including: length of long shoot, length of longest root, number of shoots, number of roots, number of nodes, number of aerial shoots, total biomass accumulation. A total of five replicate plates per species (*C. regelii* and *S. longipes*) and per temperature treatment were used. Pseudoreplicates were averaged for each replicated prior to plotting and statistical analysis. Data were analyzed by analysis of variance (ANOVA) with post hoc multiple comparisons analysis using Tukey’s honestly significant difference and plotted in GraphPad Prism v11. Experiments were replicated twice.

### Metabolomics and Hormonomics Analysis Sample Preparation

Samples were harvested from the temperature treatment experiments, rinsed to remove any tissue culture medium, patted dry and frozen at −20 °C until analysis. Methods for metabolomics and hormonomics analysis followed previously published standard operating protocols (Giebelhaus *et al*., 2024). In brief, tissues were homogenized in microcentrifuge tubes using a disposable tissue grinder (Konte Pestle Grinder) under liquid nitrogen and extracted in 50 % methanol (Fisher Scientific, Optima Grade) with 4 % acetic acid (Sigma Aldrich, LC-MS Grade) with 30 s of vortexing followed by 15 minutes sonication in ice water. Samples were centrifuged (3 min, 13, 000 rpm) and supernatant diluted 5X with water. The diluted sample was centrifuge filtered (Millipore Ultra Free MC, 0.22 µm, 1 min @ 13,000 rpm) prior to injection. Extraction solvent and water blanks were included, as well as a mixed hormonomics sample, for compound identification and quality control (Giebelhaus *et al*., 2024).

### Liquid Chromatography and Mass Spectrometry

Tissue extracts (10 μL) were separated by reverse phase chromatography (Phenomenex Kinetex EVO C18 column (2.1 x 150 mm, 17. µm Torrance, CA, USA) using a Thermo Scientific Vanquish ultra performance liquid chromatography (UHPLC) with column oven temperature of 30 °C and a flow rate of 0.250 mL/min. A gradient of A: 0.1 % aqueous formic acid B: acetonitrile was used, 0 – 25 min, 95:5 – 5:95, 25 min – 30 min, 5:95 – 95:5 with an equilibration time of 5 min between runs. Features were detected by a Q-Exactive hybrid quadrupole-orbitrap mass spectrometer with heated electrospray ionization (HESI) ionization (Thermo Scientific); settings were as follows: spray voltage: 7kV; sheath gas pressure: 48; auxillary gas flow rate: 11; sweep gas flow rate:1; capillary temperature: 250 °C; auxiliary gas heater temperature: 300 °C, S-lens RF level: 50.0, scan mode - Full MS – SIM, resolution: 70,000; polarity: positive; automatic gain control (AGC); injection time (IT): 200 ms; scan range: 100 – 1200 *m/z*; AGC target: 1E06; loop count: 5; auxillary gas heater temp: 300 °C; dynamic exclusion: spray stabilization and collision gas in the C-trap.

### Data Processing and Statistical Analysis

Raw data files were exported and processed in MZmine 3.9 (RRID:SCR_012040, Pluskal *et al*., 2010; Schmid *et al*., 2023). Mass detection used presets of MS1, all scan types and with noise level (baseline peaks) established at 3E05, a retention time window of 1-19 min and mass range of 100 – 2000 m/z. The mass list was filtered and ADAP chromatogram algorithm use to generate the extracted ion chromatograms (EIC). For deconvolution, the group intensity threshold was set to three times noise level (9E05), and the minimum number of consecutive scans set to 5. Minimum absolute height was set at 7 times noise level (2E06), with a scan to scan accuracy of 0.002 m/z or 10 ppm. Savitsky Golay smoothing was perfomred using a window of 7 scans. Chromatograph resolving was completed using the ADAP Chromatogram builder module, using the local minimum resolver with a chromatographic threshold of 90 %, minimum search range of 0.05 min, minimum relative height of 0 %, minimum absolute hiehgt of 2E06, minimum peak top/edge ratio of 1.7, minimum number of data points as a group set to 5, and a peak duriation range of 0.8 min. Settings were chosen in agreement with suggested settings for the Orbitrap detector for mzmine3. Features were then aligned across samples using join aligner with settings: m/z tolerance 0.001 or 5 ppm, weight for m/z 3, retention time tolerance 0.1 min and weight for retention time 1. Gap filling of the aligned feature list was completed using peak finder, and was applied with settings matching those used for join aligner with minimum data points 3, retention time tolerance 0.1 min and internsity tolerance 20%. Features were initially annotated by retention time and accurate mass (precursor mass) matching to the HormonomicsDB database with a 25 % retention time tolerance and mass difference of 0.02 m/z. Because the database includes both predicted and observed retention times, where retention time differed, all annotated features were compared to expected m/z and kept if within 5 ppm error regardless of matched retention time. The feature list of both annotated and unannotated features were exported with retention time, m/z ratio, and peak area to Microsoft Excel 365 (RRID:SCR_012040) for curation and further analysis.

For data curation, detected features were compared to solvent blanks, and removed if they were also found in both samples and blanks. Features were normalized to mass dry weight (DW) used during extractions. Features were then imported to Metabolanalyst 6.0 (RRID:SCR_015539) and processed using the metadata and one-factor modules. During import, data were transformed using a square root transformation, and scaled using Pareto scaling prior to statistical analysis. For untargeted metabolomic analysis, a principal component analysis was performed, followed by a two-way ANOVA using an adjusted *p*-value cutoff of 0.05 and with false discovery rate for multiple testing correction. Significant features from this analysis were visualized as a heat map. The heat map was prepared from the normalized feature list, where data were log2(*x*+1) transformed and mean-centered, then clustered by both sample and mass features using a Euclidean distance matrix and Ward’s D agglomerative clustering in Genesis 1.8.1 (RRID:SCR_015775). Annotated features were analyzed by two-way ANOVA with post hoc multiple comparisons analysis using Tukey’s honestly significant difference and plotted using GraphPad Prism v.10.

## Results

### Field Collection and Species Identification

*The permafrost anomaly itself had no signs of animal activity or otherwise that would increase in nutrient availability leading to increased plant growth. It was warm to the touch which was initially what brought our interest to the site. We hypothesize that this* may be the result of either permafrost thaw or geological activity. Field surveys identified a total of ten plant species of vascular plants (excluding grasses and sedges) were present at or adjacent to the permafrost anomaly including: Draba subcapitata Simmons, Stellaria longipes, Papaver cornwallisense D.Löve, Saxifraga cernua L., Micranthes nivalis (L.) Small, Saxifraga cespitosa L., Saxifraga oppositifolia L., Salix arctica Pall., Cochlearia groendlandica L. and Cerastium regelii. (Figure 1). Generally, the species found in the centre of the anomaly were herbaceous species, while those in the surrounding areas had a higher proportion of woody and shrubby species (Figure 1A). Stellaria longipes (Figure 1C & E) and C. regelii (Figure 1B & F) were selected for further investigation and comparison due to their high population density at the anomaly and their close phylogenetic relationship (Figure S1).

### In Vitro Germplasm Collection

Vegetative tissues of *C. regelii* and *S. longipes* were successfully established in culture. Field collected tissues were plated in small tufts on MSO media and were maintained in perpetual culture prior to conducting experiments. Both species form low growing mats across the surface of the media in these conditions. We found that maintenance in petri dishes was more successful than culture boxes as this helped to minimize contamination in *S. longipes* cultures, which appeared to be due to the presence of an endophyte which shifted to external growth from plant tissues when cultured in magenta boxes.

### Phenology

*Cerastium regelii* (Figure 2A) showed a more significant shift in phenological patterns than *S. longipes* (Figure 2B). Data from 1850-1950 shows a bimodal distribution in proportion of observations in bloom with a smaller peak in July followed by a larger maxima in September (Figure 2A, purple). The most recent observations from 2011 – 2019 show a shift towards a single slightly skewed peak in July with the intervening decades representing a gradual transition between the two distributions (Figure 2A, green). In contrast the distributions for *S. longipes* (Figure 2B) do not have a clear shift in phenological distributions over time, though there are a greater proportion of observation with flowers in later months in the year in 2011-2022 time frame compared to 1850-1950. Note that *C. regelii* observations end in 2019 while *S. longipes* end in 2022 due to limitations in the number of observations recorded in the database.

**Figure 2.**
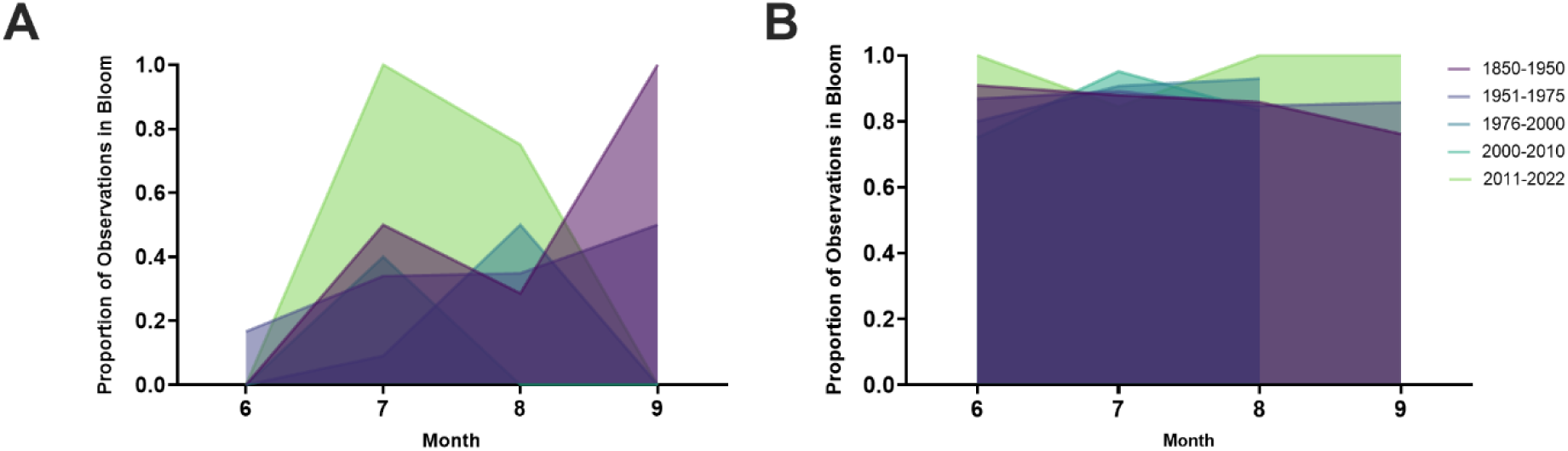
Phenological shifts in flowering time for (A) Cerastium regelii and (B) Stellaria longipes from historic observations in the Territories displayed as proportion of observations in bloom per month. More recent observations are lighter in colour.

### Ecological Niche Modeling

A total of 2,471 occurrence points for *S. longipes* and 79 for *C. regelii* were included in the final models. Differences in future ecological niche suitability in the Canadian High Arctic showed a stark contrast between the two species. The models generated predict little change in niche suitability for *S. longipes* between current and future climate scenarios (Figure 3B) but a significant increase in future niche suitability for *C. regelii* (Figure 3C). National models align with current distributions of *C. regelii* with suitable habitat occurring only in the High Arctic with no range expansion predicted (Figure S2). *Stellaria longipes* national models (Figure S3) again show a more moderated response with marginal improvements in niche suitability in Northern and alpine regions. Under Shared Socioeconomic Pathway (SSP) 126, Resolute (Qausuittuq) is predicted to experience an ∼5 °C increase in mean annual temperature, with this increasing to almost 10°C under SSP 585 (Table 1).

**Figure 3.**
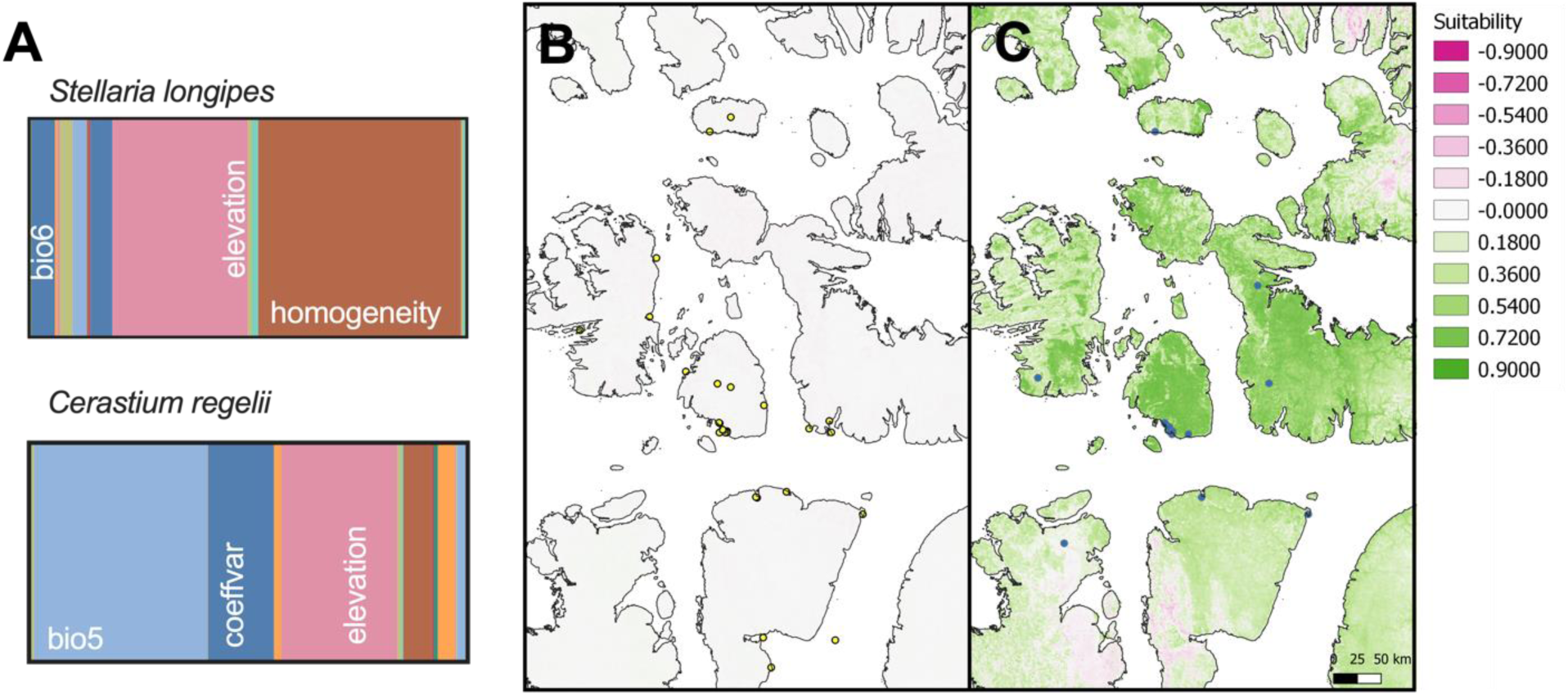
(A) Variable contribution to ecological niche models for Stellaria longipes and Cerastium regelii and predicted change in suitability based on maxent models for SSP 126 projected to 2080 for the region around Cornwallis Island for (B) Stellaria longipes, yellow and (C) Cerastium regelii, blue where points indicate occurrence records included in the model, green and increase in predicted niche suitability, white no change and pink a decrease in predicted niche suitability as compared to current range predictions.

**Table 1.** Average annual temperature (BIO1) of Cornwallis Island (−96.554589, 75.64527, −93.1-1039, 74.515367) sampled at 1,000 random points under historic and mean future climate scenarios.

|  | Annual mean ( °C)<sup>2</sup> | Annual delta (°C) |
| --- | --- | --- |
| Historic | -17.19 |  |
| SSP585<sup>1</sup> | -7.50 | 9.69 |
| SSP126<sup>1</sup> | -12.11 | 5.08 |

<sup>1</sup>Average of CNRM, MRI & MIROC projections
<sup>2</sup>Calculated with no data values removed

Variable contributions to models for *C. regelii* and *S. longipes* differed strongly as reflected in the magnitude of the change in niche suitability under future climates. The largest contributor to models for *C. regelii* was a bioclimatic variable, bio5 (39%, maximum temperature of the warmest month, a bioclimatic variable), followed by elevation and coefficient of variation of habitat heterogeneity (Figure 3A). Habitat homogeneity and elevation were the two largest contributors to the model for *S. longipes*, while the highest contributing bioclimatic variable was bio6 (minimum temperature of the coldest month), with a contribution of only ∼ 5% (Table S3).

### In Vitro Experimental Warming

Field grown tissues grown under three temperatures 12, 24 or 28 °C of *Stellaria longipes* had a significant maxima in growth across all parameters determined at 24 °C (Figure 4). While *C. regelii* showed a similar and significant pattern in growth for root growth and shoot height, the number of shoots showed a linear increase as temperature increased, with the highest number of shoots produced at 28 °C (Figure 4C).

**Figure 4.**
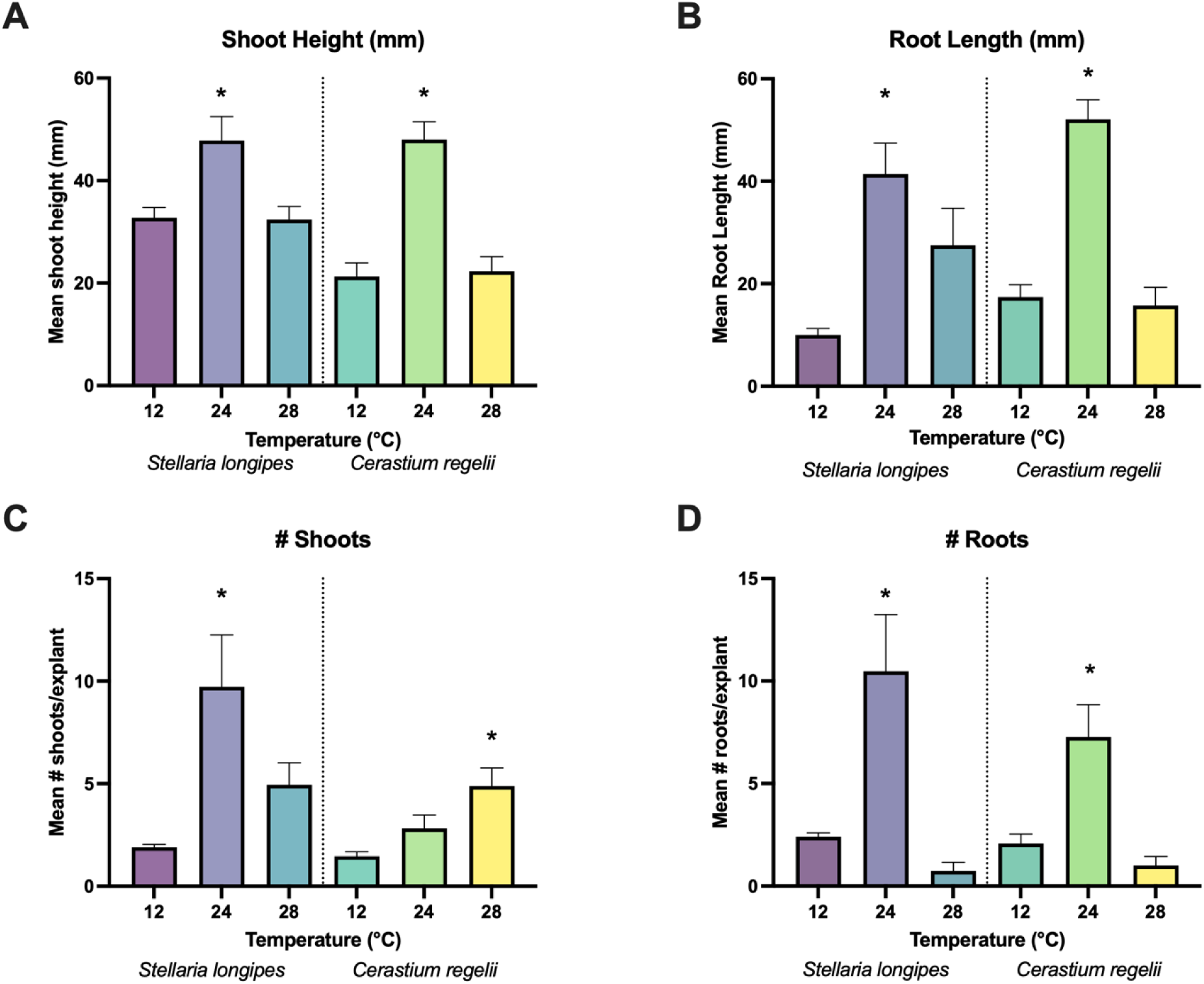
Changes in selected growth parameters for Cerastium regelii and Stellaria longipes after 3 weeks at 12, 24, or 28 °C. Both species had maximal increases to shoot height (A), root length (B), and number of roots (D) at 24 °C. S. longipes also had maximal increase to number of shoots at this temperature, but C. regelii instead showed increasing number of shoots with increasing temperature, with maximum number of shoots observed at 28 °C. Data were analysed with one-way ANOVA with Tukey’s test for multiple comparisons for each species, asterisks represent significant differences (α = 0.05) within a parameter and species. Error bars represent standard error.

### Metabolomics and Hormonomics

Following UPLC-MS analysis of plant extracts, a total of 345 mass features were detected, and 11 of these were annotated as hormones, representing predominantly metabolites from the cytokinin or tryptophan metabolism pathways (Table S4). From the pool of both annotated and unannotated mass features, a two-way ANOVA revealed 274 mass features that were differentially abundant between species and across increasing temperature (Figure 5A). 18 of the features identified via ANOVA were unique to species and 14 to temperature. 74 mass features were significantly different when either species or temperature were considered, 71 of which had a significant interaction between temperature and species. Principal component analysis showed that the greatest variation within the dataset was due to species, as shown in principal component 1 representing 28 % of variation (Figure 5B). However, PC2 and PC3 (not shown) collectively separated samples by temperature, representing a total of ∼36 % of all variation.

**Figure 5.**
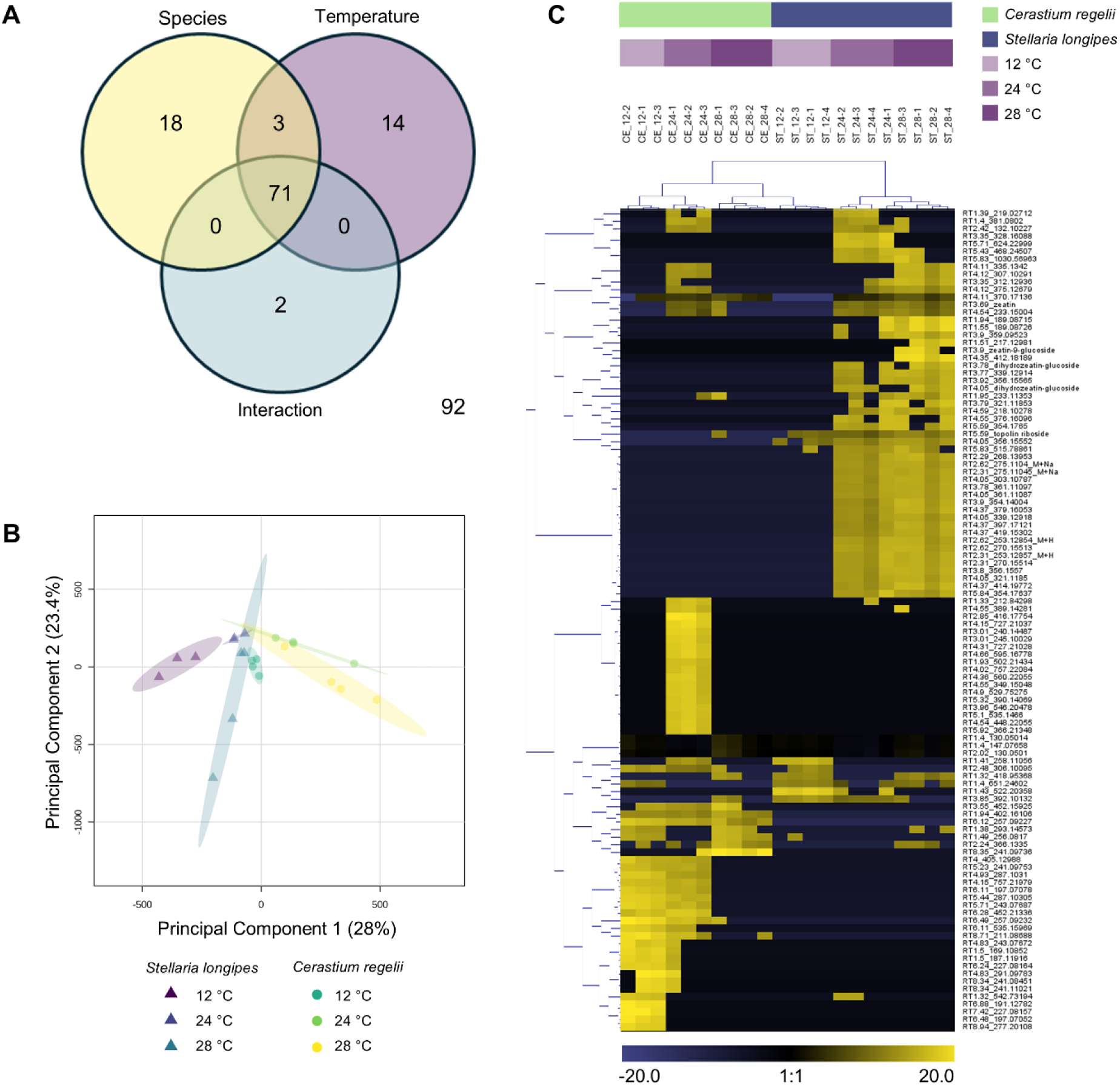
Untargeted metabolomic analysis of Stellaria longipes and Cerastium regelii extracts after growth at 12, 24, or 28 °C for 3 weeks. A two-way ANOVA shows unique mass features related to species or temperature, and a large proportion of mass features shared that have an interaction (A). Principal component analysis (B) separates samples by species along PC1, representing the most variation within the dataset. A heat map prepared using all significant mass features clustered using Euclidean distance and Ward’s D method (C) highlights strong differences between species and temperature. A large group of mass features are mostly unique to S. longipes grown at 24 or 28 °C, including some mass features annotated as cytokinins involved in shoot growth and elongation.

Hierarchical clustering produced relatively distinct clusters for each condition tested, with samples of a specific species and temperature being closely related (Figure 5C). *C. regelii* clustered strongly together, along with *S. longipes* grown at low temperatures. A second large cluster included *S. longipes* grown at 24 and 28 °C, with some intermixing between treatments, indicating *S. longipes* significantly modifies metabolism in response to increased temperature. Notably, this includes cytokinins topolin riboside, dihydrozeatin glucoside and zeatin-9-glucoside. These cytokinins were not detected in *C. regelii* at 12 or 28 °C, only at 24 °C.

### Cytokinin Metabolism

Most annotated features were related to cytokinin biosynthesis. Zeatin increased with increasing temperature in both *C. regelii* and *S. longipes* (Figure 6), but the normalized abundance was similar between species at each temperature. In contrast, dihydrozeatin, dihydrozeatin glucoside, and zeatin-9-glucoside were not detected in *C. regelii*, but significantly increased (*α* < 0.05) with temperature in *S. longipes*, with the highest abundance at 28 °C, suggesting that *C. regelii* and *S. longipes* utilize different signalling pathways to adapt to high temperature. Zeatin-7-glucoside abundance in *C. regelii* mirrored growth patterns observed for shoot height, with maximum abundance at 24 °C representing a significant increase (*p* = 0.033), but was absent in *S. longipes* except at 28 °C. Topolin riboside increased with increasing temperature for both species, but was in greater abundance *S. longipes* than *C. regelii*, to a significant extent (*p* = 0.0046).

**Figure 6.**
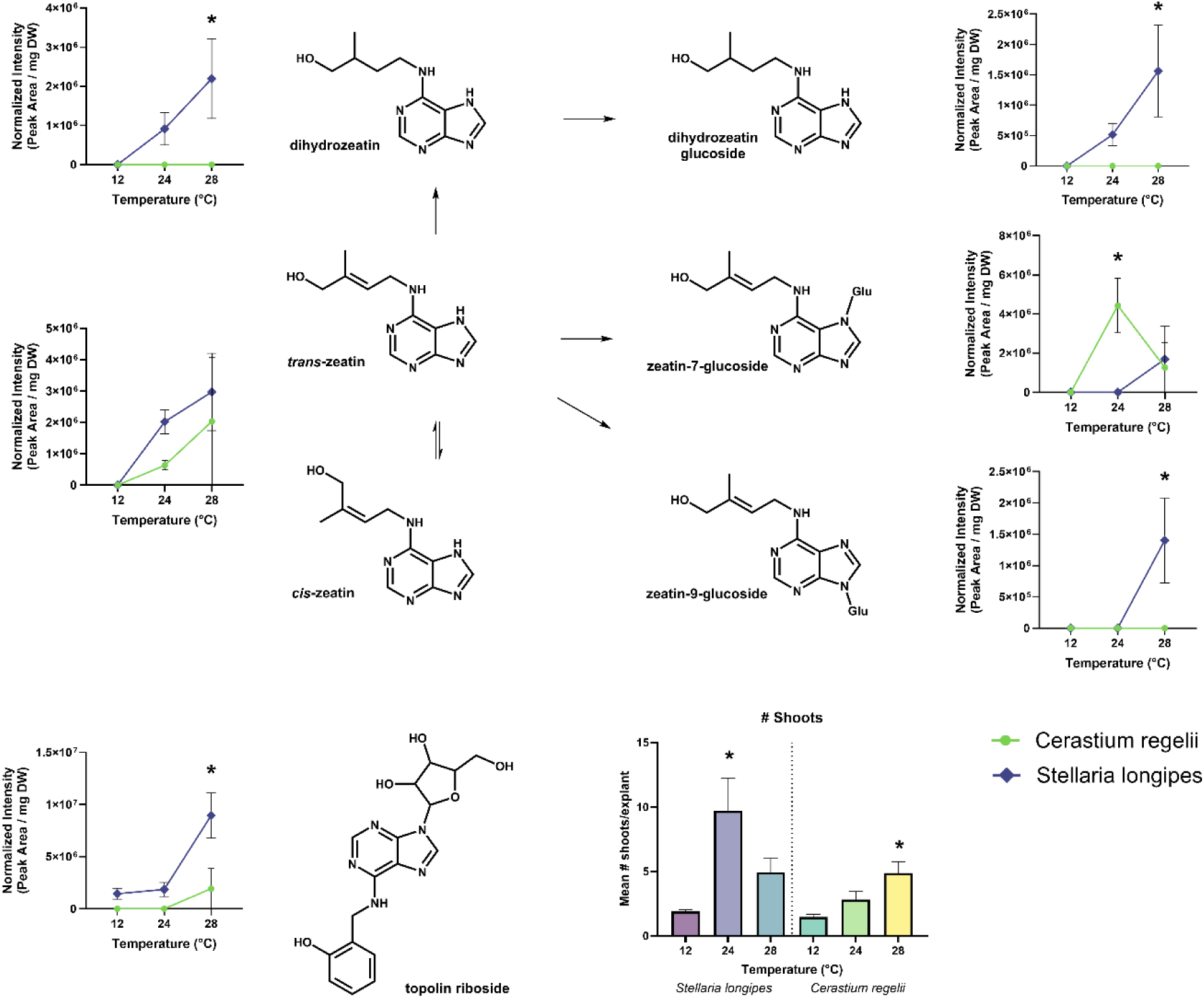
Effects of temperature on cytokinin metabolism for zeatin, dihydrozeatin, dihydrozeatin glucoside, zeatin-7-glucoside, zeatin-9-glucoside, and topolin-riboside in Stellaria longipes and Cerastium regelii. For clarity, only the structures for trans-zeatin-7-glucoside, trans-zeatin-9-glucoside and ortho-topolin riboside are depicted, but this analysis did not differentiate between isomers. Shoot growth patterns are shown below for reference. Error bars represent standard error.

## Discussion

Following a field survey of a permafrost anomaly in Resolute (Qausuittuq), Nunavut, we established cultures of *S. longipes* and *C. regelii* for *in vitro* studies, along with field-collected seeds. We performed experimental warming to test the effects of rising temperature in comparison to field observation and predicted ecological niche modelling for both *S. longipes* and *C. regelii,* focusing on impacts to reproductive and vegetative growth. We hypothesized that between these related species, as a more generalist species with a greater distribution, *S. longipes* would be more resilient to temperature increases than *C. regelii*, as evidenced by increased growth and a modified phytohormone profile.

Field observations of the permafrost anomaly showed that for the plant community established there, an increase in temperature shifted species composition towards herbaceous plants including *S. longipes* and *C. regelii*, suggesting that increased warming created a more suitable habitat for both species in these conditions. As species in this area are not currently in competition with heavy shrub cover yet, our observations may provide insight into early succession stages prior to shrubification associated with arctic greening, following tundra fires, or following browning events with high mortality (Box *et al*., 2019; Chen *et al*., 2024; Phoenix *et al*., 2025). Increased herbification with warming was largely supported by ecological niche modeling for these species. Under SSP126 Resolute (Qausuittuq) is predicted to experience an average temperature increase of ∼5 °C. In these models *S. longipes* shows very little change in habitat suitability across its current range, with only a slight Northward expansion. In contrast, habitat suitability in current regions for *C. regelii* is predicted to significantly increase, although its range will remain similar.

Within these models, the main contributors of environmental and bioclimatic variables to model outcomes were different between species. Variables relating to temperature did not strongly contribute to niche suitability, while habitat homogeneity and elevation was highly predictive of niche suitability for *S. longipes*. In contrast, the main contributor to niche suitability for *C. regelii* was the maximum temperature of the warmest month, highlighting a temperature-dependent relationship of niche suitability and range expansion for *C. regelii* not predicted for *S.* longipes. This is consistent with our hypothesis that *S. longipes*, as a generalist, would be less sensitive to temperature change in moderate scenarios such as those explored here. This also suggests that within these scenarios, increased warming may have a positive, rather than negative influence on *C. regelii.* Phenology is the measure of the biological life cycles over an annual cycle, and includes important events such as timing of bud break, flowering, fruiting, and dormancy (Morales-Castilla *et al*., 2024). As these events are closely linked to environmental conditions, phenology has been increasingly used as an indicator of climate change, but it is important to investigate this across a broad range of species and climes as both phylogenetics and environment play a role in phenological shifts (Diepstraten *et al*., 2018; Morales-Castilla *et al*., 2024). In arctic climes, the effects of climate change are accelerated, and early loss of sea ice, increased temperature and radiation, and a longer growing period are contributing to changes in flowering time, fruit and bud set, along with pollinator emergence (Stewart *et al*., 2018; Box *et al*., 2019). We evaluated the phenology of *S. longipes* and *C. regelii* over the past 170 years to evaluate how warming temperatures affected the flowering period. In most species, the flowering period is compressed, with phenological shifts towards earlier flowering times such as in *Dryas integrifolia* Vahl where early flowering time has been linked to monthly temperature just prior to flowering (Panchen & Root, 2015). We found that neither *S. longipes* nor *C. regelii* flowered earlier. *S. longipes* experienced no change in flowering time. The number of plants in bloom observed increased with time, but may reflect increased access to the area or similar observational bias during this time period. *C. regelii* did undergo contraction of the flowering period from a bimodal distribution with two blooming periods in July and September, to a unimodal distribution over a shorter flowering period, but this also resulted in a general shift to later blooming time in late July to early August. Based on niche suitability modelling and historical phenology, *C. regelii* exhibits a more pronounced change in characteristics due to temperature than *S. longipes,* but contrary to our hypothesis, this may actually improve abundance and suitability for this species within this region.

Based on the modelled results, we followed up with *in vitro* experiments to test the effects of temperature increases on growth and phytohormone profiles of these two species experimentally. We grew cultured *S. longipes* and *C. regelii* at 12 °C (a typical summer temperature in the High Arctic), 24 °C (a typical summer temperature in the southern-most latitudes where *S. longipes* is found), and 28 °C to model increased warming for these areas. An increase in temperature from 12 to 24 °C, significantly increased the number of shoots and roots produced in *S. longipes*, accompanied by increases in shoot and root length. Further increases of temperature to 28 °C had the opposite effect, reducing these parameters to values observed at 12 °C. These patterns were also observed in *C. regelii* for the number and length of roots, and the length of shoots, but the number of shoots increased linearly with increasing temperature. G rowth trends were similar between these two species for most conditions, which may reflect on their similar abundance and suitability at the temperatures observed in the permafrost anomaly. An opposite trend in *C. regelii* associated with number of shoots shows that temperature does affect some aspects of growth differently for *C. regelii* and *S. longipes*, which may have downstream implications for their individual adaptability. *C. regelii* shows consistent increases in the number of shoots it produces even at the highest temperature included in this study, while *S. longipes* shows reduced number of shoots at these temperatures.

Given that plant growth is regulated by cross-talk between a growing number of phytohormone classes, we theorized that differences in growth, such as in the number of shoots, could be resolved by exploring differences in phytohormone production (Mughal *et al*., 2024; Banoriya *et al*., 2024). We used metabolomics to broadly explore changes to metabolomic profile and phytohormone production. The primary driver of variability in metabolites was species, but following hierarchical analysis, we found that warming temperature has a strong influence on both species. We had expected to see the greatest differences in metabolites at the temperatures which were associated with the biggest increases in growth, 24 °C for *S. longipes*, and 28 °C for *C. regelii*. What we observed in our results, however, was that the largest differences for *S. longipes* was when it was warmed past the optimal temperature. Given that growth patterns tended to be self-limiting with a reduction back to normal growth patterns at 28 °C, additional changes to the metabolite profile are thus likely indicative of an early stress response. These results support the predictions generated by ecological niche suitability modeling and phenology analysis, where *S. longipes* was not predicted to change substantially in suitability for the region studied. The results for *C. regelii* are more challenging to interpret. Although the responsiveness of *C. regelii* was highest at 24 °C, we continued to see increased growth in the number of shoots beyond this temperature. Based on these results, we hypothesized that the production of specific phytohormones required for growth is differentially synthesized or regulated between these two species, and is more influenced by temperature in *C. regelii* that is reflected in niche suitability modelling.

Following the complex growth response of *C. regelii* to temperature, we explored the abundance of phytohormones annotated from our metabolomics analysis. The strong effect of experimental conditions on shoot growth and number highlights a relationship between temperature and the production of cytokinins. Cytokinins are strongly associated with shoot growth, and are known to stimulate cellular proliferation, vascular development, and biomass distribution in addition to abiotic stress responses like heat (Li *et al*., 2021). Zeatin, a major cytokinin, was similarly produced by both *S. longipes* and *C. regelii* as temperature increased. Zeatin is a major isoprenoid cytokinin produced by most plants as *cis-* or *trans-*zeatin, and is one of the bioactive forms with strong bioactivity (Sakakibara, 2006). Other cytokinin derivatives strongly differ between species. Dihydrozeatin, a bioactive zeatin derivative steadily increased in abundance in *S. longipes* as temperature increased, but this coincided with an increase in dihydrozeatin glucoside especially at 28 °C, widely considered to be an inactive form. *S. longipes* also produced detectable quantities of two other inactive zeatin forms, zeatin-7-glucoside and zeatin-9-glucoside. Collectively, the production of increased inactivated zeatin derivatives at the highest temperature is likely to have a dampening effect on growth, supporting the observed decrease in growth for *S. longipes* at this temperature. In *C. regelii,* the two isoprenoid cytokinins detected were zeatin and the inactivated zeatin-7-glucoside. Zeatin-7-glucoside was most abundant at 24 °C. At 28 °C, zeatin-7-glucoside decreased alongside a corresponding increase in zeatin, and this may help explain the increased growth observed at this temperature. Lastly, both *S. longipes* and *C. regelii* produced an abundance of topolin riboside, an aromatic cytokinin found in a limited number of plant species (Zhao *et al*., 2024). It was increased in abundance at 28 °C, although in significantly greater abundance in *S. longipes*, and may suggest an alternative cytokinin production pathway under heat stress. These results show that differences in these species as they react to temperature increases are governed by alternative hormone signaling pathways. Further investigations are needed to determine the influence of temperature on other hormone classes, and given the differences in cytokinin abundance, what cross-talk may exist between them.

Collectively, our results show two strikingly different outcomes for these two herbaceous species of the Arctic tundra. *S. longipes* shows very little influence of temperature on its niche suitability or phenology, and may maintain vigor and robustness under the moderate future climate scenarios explored here, supporting our hypothesis that as a generalist species it would be robust to change at increased temperatures. In direct contrast, although *C. regelii* is more affected by temperature, but for the conditions evaluated in our study is generally positively influenced with warming, which is surprising given that it is more specialized with a smaller distribution and predicted lower tolerance to change. Our *in vitro* investigations supported these predictions, showcasing that *S. longipes* is self-limiting with temperature changes, while increased temperatures tend to favor increased growth in *C. regelii.* Despite steady increases in cytokinin production that typically induce increased growth, *S. longipes* is self-limiting within the temperature ranges evaluated. Yet our work highlights the importance of studying climate change in the Arctic not only as a function of ecosystems and biomes, but also as a function of species, location, and metabolism. It is clear that even related species such as *S. longipes* and *C. regelii* can have different and unexpected responses to both moderate and extreme temperature increases as explored here. We also show the benefit and importance of using techniques like metabolomics to investigate these questions, helping us to evaluate how warming temperatures of the future will impact growth patterns and other aspects of plant biology for a variety of species with different responses to climate change. Our study also expands our knowledge and understanding of *C. regelii*, an understudied and important herbaceous plant in the Canadian High Arctic.

## Supporting information

Full Supporting Information

## Acknowledgements

Funding support was provided by the Alpine Club of Canada Environment Grant, the Explorer’s Club Mamont Scholar Grant and the Natural Sciences and Engineering Council of Canada to Dr. Erland. Logistical support was provided by the Polar Continental Shelf Program (PCSP) of the National Research Council of Canada (Project: 63319 – “ARCTIC Change - Arctic Research and Conservation Team Investigating Climate Change”). Guidance for this project was provided by staff and researchers at the PCSP facility in Resolute (Qausuittuq). We are grateful to Brandy Iqaluk, our guide in Resolute (Qausuittuq). We would also like to acknowledge undergraduate students at the University of the Fraser Valley who contributed to visual assessment of the phenology results Jiadong Li, Sebastian Molina, Elisha Marks and Oscar Caunce. Ryland Giebelhaus who assisted in maintenance of the initial cultures, and Sarah Lyons at Supra RnD Labs who assisted in acquisition of the metabolomics data. We are grateful to Paul Sokoloff at the Canadian Museum of Nature for his guidance in, and identification of the botanical vouchers.

## Author Contributions

L.A.E Erland performed experimental work, data analysis, writing, and editing of manuscript. S.L. Lane performed data analysis for metabolomics experiments, writing, and editing of manuscript.

## Data Availability

The metabolomic data from this study is available at the NIH Common Fund’s National Metabolomics Data Repository (NMDR) website, the Metabolomics Workbench, https://www.metabolomicsworkbench.org where it has been assigned Study ID ST003834. The data can be accessed directly via its Project DOI: http://dx.doi.org/10.21228/M8625

## Supporting Information

**Figure S1.** Phylogenetic tree of field collections from 2019 based on DNA fingerprinting data uploaded to BOLD and generated in PhyloT, with species of interest highlighted in purple.

**Figure S2.** Predicted change in ecological niche suitability for *Cerastium regelii* under current and future climates.

**Figure S3.** Predicted change in ecological niche suitability for *Stellaria longipes* under current and future climates.

**Table S1**. Summary of historic climate variables assessment for inclusion in model.

**Table S2.** Model performance statistics for the null and empirical models generated for ecological niche modelling of *Cerastium regelii* and *Stellaria longipes*.

**Table S3.** Maxent ecological niche model performance characteristics and variable contribution for variables retained in the model for *Cerastium regelii* and *Stellaria longipes*.

**Table S4.** Annotations for mass features listed by mass-to-charge ratio and retention time using indicated method.

