## Supplementary material for "Differential temperature adaptation mechanisms in the High Arctic-adapted *Cerastium regelii* Ostenf. and the widespread *Stellaria longipes* Goldie": Full Supporting Information

The following Supporting Information is available for this article:

**Fig. S1** Phylogenetic tree of field collections from 2019 based on DNA fingerprinting data uploaded to BOLD and generated in PhyloT, with species of interest highlighted in purple.

**Fig. S2** Predicted change in ecological niche suitability for *Cerastium regelii* under current and future climates.

**Fig. S3** Predicted change in ecological niche suitability for *Stellaria longipes* under current and future climates.

**Table S1** Summary of historic climate variables assessment for inclusion in model.

**Table S2** Model performance statistics for the null and empirical models generated for ecological niche modelling of *Cerastium regelii* and *Stellaria longipes*.

**Table S3** Maxent ecological niche model performance characteristics and variable contribution for variables retained in the model for *Cerastium regelii* and *Stellaria longipes.*

**Table S4** Annotations for mass features listed by mass-to-charge ratio and retention time using indicated method.

**Fig. S1** Phylogenetic tree of field collections from 2019 based on DNA fingerprinting data uploaded to BOLD and generated in PhyloT. Species of interest are highlighted in purple.

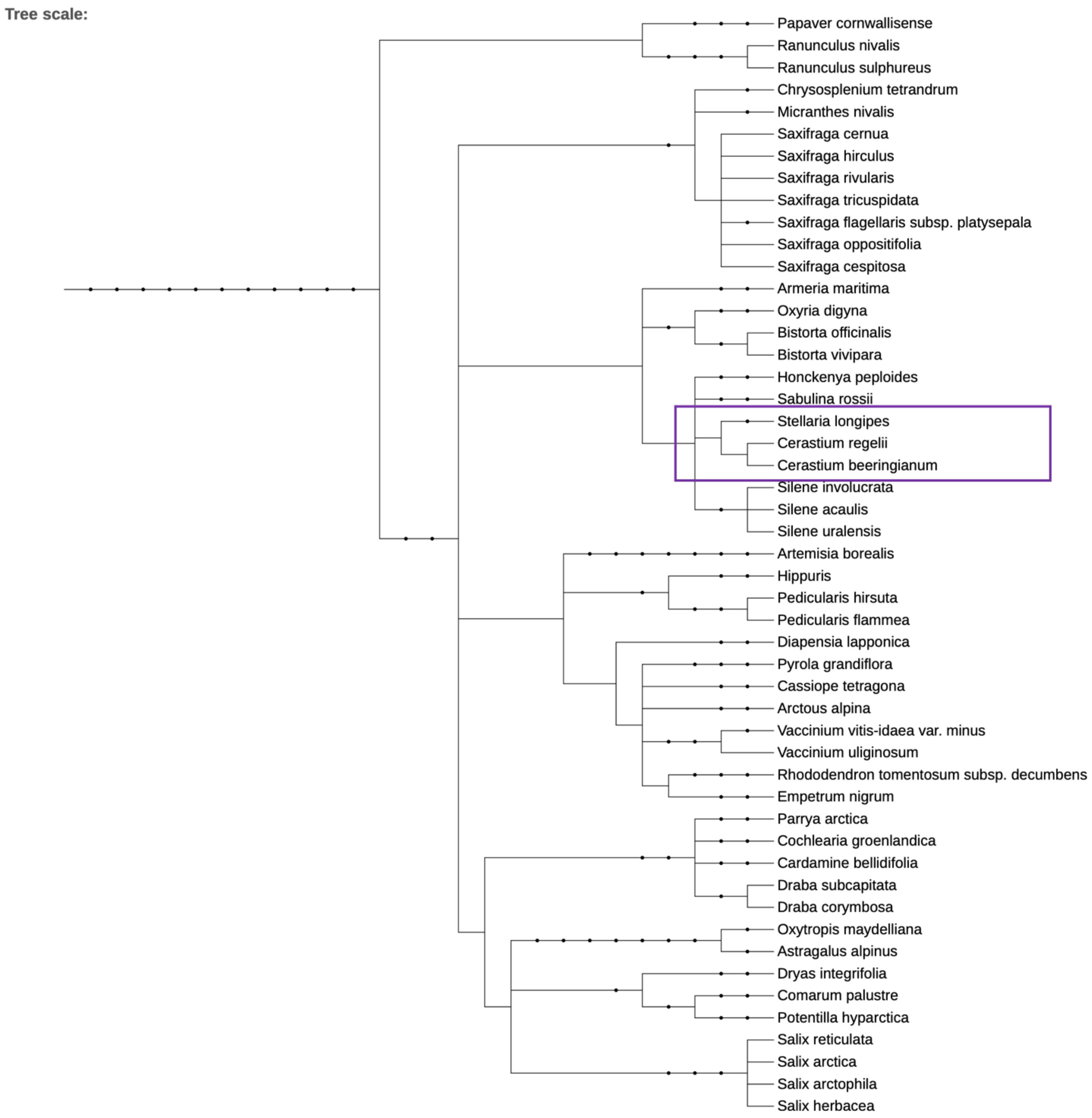

**Fig. S2** Predicted change in ecological niche suitability for *Cerastium regelii* under current and future climates. Inset shows a closer view of the Canadian High Arctic where *C. regelii* is known to occur.

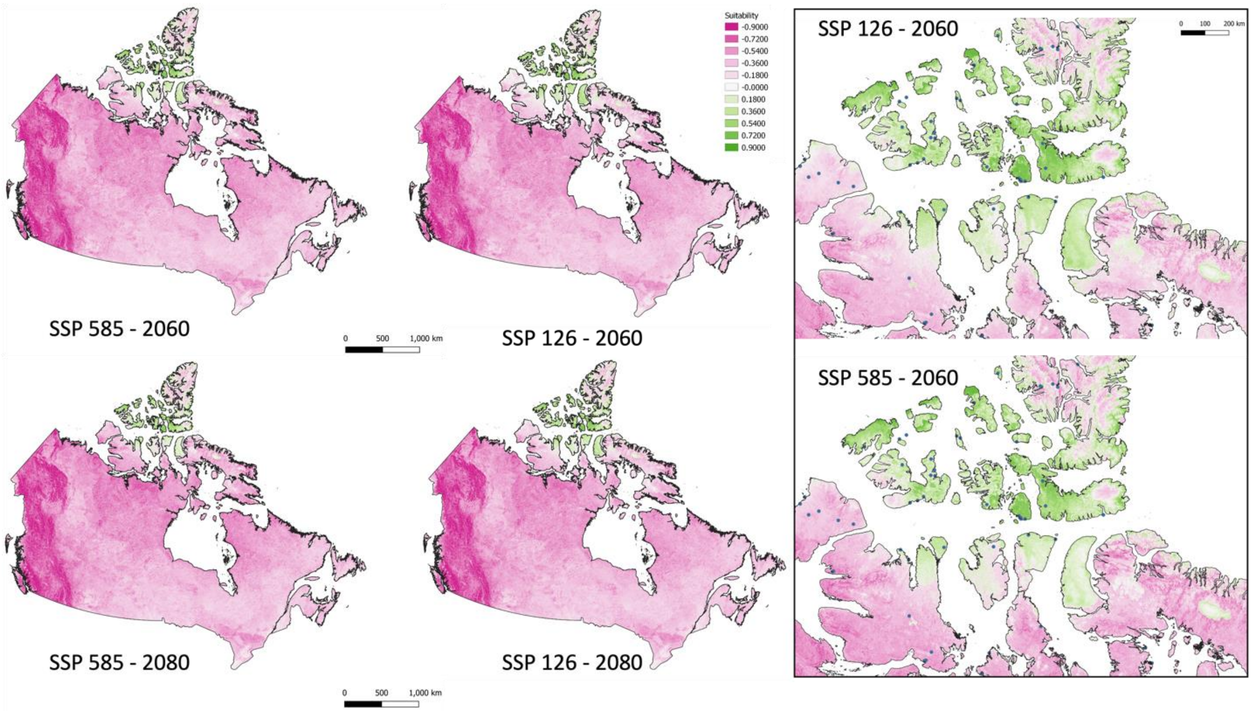

**Fig. S3** Predicted change in ecological niche suitability for *Stellaria longipes* under current and future climates

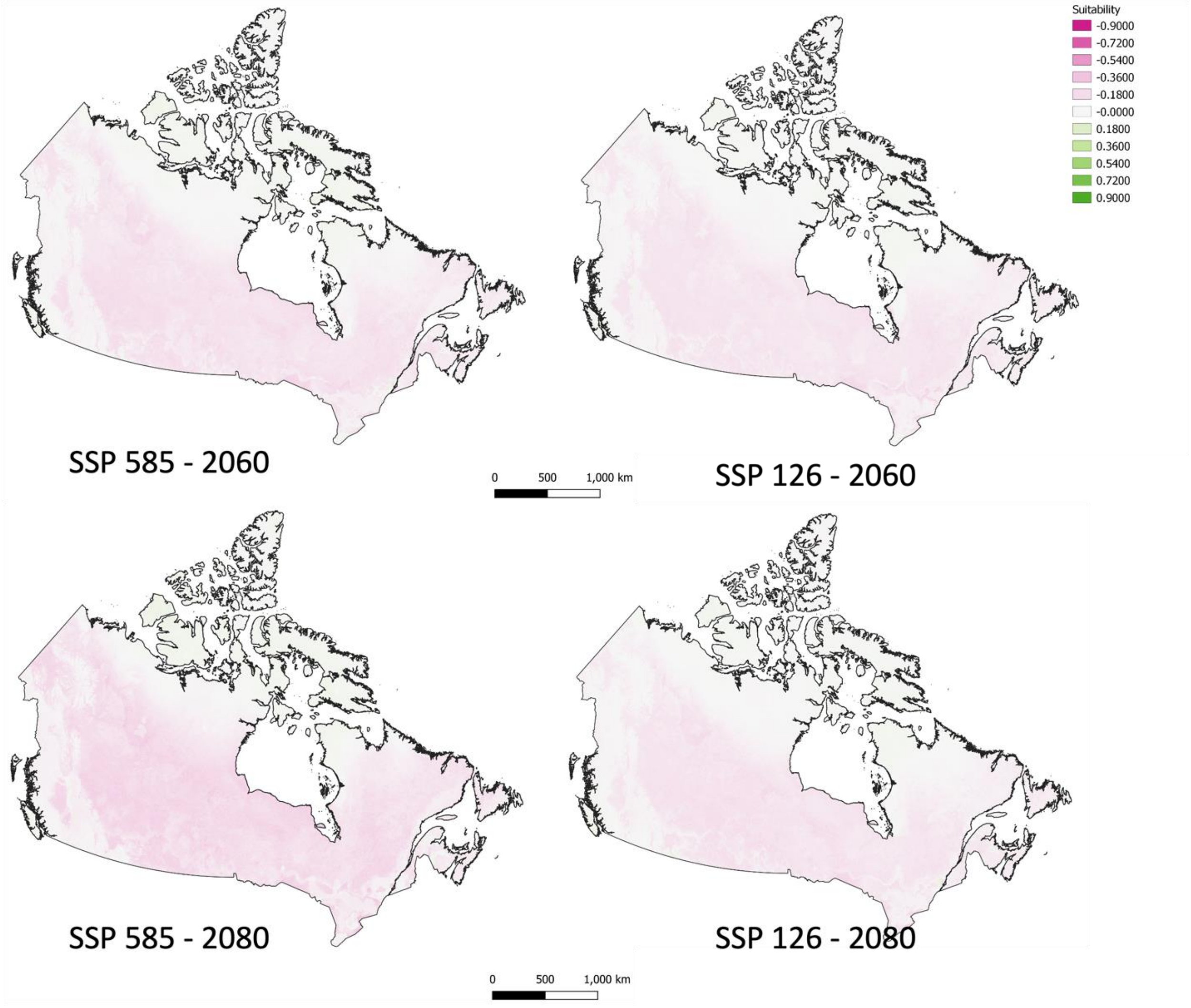

**Table S1** Summary of historic climate variables assessment for inclusion in the model

| Variable | Variable Shorthand | Source/Type |
|---|---|---|
| Annual Mean Temperature (°C) | bio1 | WorldClim |
| Maximum Temperature Warmest Month (°C) | bio5 | WorldClim |
| Minimum Temperature Coldest Month (°C) | bio6 | WorldClim |
| Mean Temperature of Wettest Quarter (°C) | bio8 | WorldClim |
| Mean Temperature of Driest Quarter (°C) | bio9 | WorldClim |
| Annual Precipitation (mm) | bio12 | WorldClim |
| Precipitation Wettest Quarter (mm) | bio16 | WorldClim |
| Precipitation Driest Quarter (mm) | bio17 | WorldClim |
| Precipitation Warmest Quarter (mm) | bio18 | WorldClim |
| Precipitation Coldest Quarter (mm) | bio19 | WorldClim |
| Aspect Sine | aspectsine | topography |
| Aspect Eastness | eastness | topography |
| Aspect Northness | northness | topography |
| Geomorphological landform (Shannon Index) | geom | topography |
| Slope | slope | topography |
| Cultivated & Managed Vegetation (% consensus prevalence) | cultivatedmanagedveg | landcover DIScover |
| Regularly Flooded Land (% consensus prevalence) | regflooded | landcover DIScover |
| Snow & Ice Cover (% consensus prevalence) | snowice | landcover DIScover |
| Homogeneity | homogeneity | habitat heterogeneity |
| Entropy | entropy | habitat heterogeneity |
| Dissimilarity | dissimilarity | habitat heterogeneity |
| Evenness | evenness | habitat heterogeneity |
| Co-efficient of Variation | coeffvar | habitat heterogeneity |
| Soil Organic Carbon Densities (hg/m3) | ocd | SoilGrids |
| Soil Organic Carbon Stock (t/ha) | ocs | SoilGrids |
| Soil Organic Carbon Content (dg/kg) | soc | SoilGrids |
| Silt (g/kg) | silt | SoilGrids |
| Sand (g/kg) | sand | SoilGrids |
| Nitrogen (cg/kg) | nitrogen | SoilGrids |

| | | |
|---|---|---|
| Clay (g/kg) | clay | SoilGrids |
| Bulk Density of Fine Earth Fraction (cg/cm3) | bdod | SoilGrids |
| Volumetric fraction of coarse fragments > 2mm (cm2/dm3 / vol%) | cfvo | SoilGrids |
| Cation exchange capacity of the soil (mmol(c)/kg) | cec | SoilGrids |
| Soil pH (pH x 10) | phh2o | SoilGrids |

**Table S2** Model performance statistics for the null and empirical models generated for ecological niche modelling of Cerastium regelii and Stellaria longipes.

| **Statistic** | **AUC** | | **Minimum Training Presence Omission Rate** | | **10 Percent Omission Rate** | |
|---|---|---|---|---|---|---|
| | null | empirical | null | empirical | null | empirical |
| *Cerastium regelii* RM = 4, Features = LQH | | | | | | |
| mean | 0.558 | 0.804 | 0.000 | 0.000 | 0.105 | 0.103 |
| standard deviation | 0.032 | 0.054 | 0.001 | 0.000 | 0.030 | 0.091 |
| zscore | 7.604 | | -0.118 | | -0.060 | |
| pvalue | <0.0001 | | 0.453 | | 0.476 | |
| *Stellaria longipes* RM = 5, Features = LQH | | | | | | |
| mean | 0.797 | 0.865 | 0.000 | 0.000 | 0.034 | 0.083 |
| standard deviation | 0.026 | 0.039 | 0.002 | 0.000 | 0.019 | 0.052 |
| z-score | 2.654 | | -0.188 | | 2.526 | |
| p-value | 0.004 | | 0.425 | | 0.994 | |

**Table S3** Maxent ecological niche model performance characteristics and variable contribution for variables retained in the model for Cerastium regelii and Stellaria longipes.

| | *Stellaria longipes* | | *Cerastium regelii* | |
|---|---|---|---|---|
| **Variable** | **Percent Contribution** | **Permutation Importance** | **Percent Contribution** | **Permutation Importance** |
| bdod | 0.0193 | 0.9225 | 1.1442 | 2.8595 |
| bio1 | 0.4998 | 0.1934 | ~~0~~ | ~~0~~ |
| bio12 | 2.8942 | 0.5572 | ~~0~~ | ~~0~~ |
| bio16 | 3.2335 | 0.6501 | 0.1314 | 0.0036 |
| bio17 | 0.5264 | 7.1554 | ~~0~~ | ~~0~~ |
| bio18 | 0.2939 | 4.4969 | ~~0~~ | ~~0~~ |
| bio19 | 0.0088 | 0.0071 | 0.0522 | 0.0044 |
| bio5 | 0.0295 | 0.0794 | 39.4382 | 42.7297 |
| bio6 | 5.3681 | 0.9594 | ~~0~~ | ~~0~~ |
| bio8 | 0.5832 | 0.917 | ~~0~~ | ~~0~~ |
| bio9 | 0.5934 | 5.0889 | 0.0527 | 0.0285 |
| cec | 0.0199 | 0 | ~~0.0716~~ | ~~0.8538~~ |
| coeffvar | 4.98 | 0 | 14.9568 | 0 |
| dissimilarity | 0.0408 | 0 | 1.6561 | 0 |
| elevation | 30.906 | 25.3389 | 26.4285 | 16.868 |
| entropy | 0.8703 | 0 | 0.9293 | 0 |
| evenness | 1.4589 | 1.1872 | 0.4248 | 15.1271 |
| geom | 0.0111 | 0.0864 | ~~0~~ | ~~0~~ |
| homogeneity | 46.0241 | 38.5921 | 6.6315 | 19.4896 |
| nitrogen | 0.1316 | 4.3129 | 0.3514 | 0.5358 |
| northness | 0.0204 | 0.6623 | ~~4.00E-04~~ | ~~0~~ |
| ocd | 0.1348 | 5.6983 | 0.8418 | 0.781 |
| ocs | ~~0~~ | ~~0~~ | 0.0064 | 0.0372 |
| openwater | 0.3542 | 0 | 3.8697 | 0.1823 |
| phh2o | ~~0~~ | ~~0~~ | 0.2547 | 0 |
| silt | ~~0~~ | ~~0~~ | 0.2982 | 0 |
| slope | 0.6206 | 1.008 | ~~0.011~~ | ~~0.0617~~ |
| snowice | 0.3013 | 1.4626 | 2.4147 | 0.0176 |
| soc | 0.0211 | 0.447 | ~~0~~ | ~~0~~ |

| | | |
|---|---|---|
| TSS Train | 0.528 | 0.777 |
| TSS test | 0.511 | 0.73 |
| TSS heldback | 0.546 | 0.81 |
| AICc | 59554.01 | 31389.84 |
| AUC | 0.782 | 0.929 |

**Table S4** Annotation of mass features. Mass features are listed by m/z and retention time in minutes. Annotations made by exact mass are within 5 ppm error of the reference mass. Annotations made by local database were matched by m/z and retention time using annotation functionalities in mzmine 3.9.

| Mass Feature | Compound Annotation | Database | Annotation Method |
|---|---|---|---|
| 138.0555_1.44 | anthranilic acid | HormonomicsDB | Exact Mass |
| 182.08172_2.06 | tyrosine | HormonomicsDB | Exact Mass |
| 203.22357_1.28 | spermine | HormonomicsDB | Exact Mass |
| 205.09718_4.29 | tryptophan | HormonomicsDB | Local Database |
| 220.11931_3.69 | zeatin | HormonomicsDB | Local Database |
| 366.17775_6.15 | N6 isopentenyladenine-7-glucoside | HormonomicsDB | Exact Mass |
| 374.14592_5.59 | Topolin riboside | HormonomicsDB | Local Database |
| 382.17213_3.9 | zeatin-9-glucoside | HormonomicsDB | Local Database |
| 384.18778_3.78 | dihydrozeatin-glucoside | HormonomicsDB | Local Database |
| 384.18778_4.05 | dihydrozeatin-glucoside | HormonomicsDB | Local Database |
| 388.16209_5.91 | N6-benzyladenine-7-glucoside | HormonomicsDB | Exact Mass |
